# Phylogenetic and Ecological Trends in Specialization: Disentangling the Drivers of Ectoparasite Host Specificity

**DOI:** 10.1101/2022.04.06.487338

**Authors:** Alexis M. Brown, Kelly A. Speer, Tiago Teixeira, Elizabeth Clare, Nancy B. Simmons, Juan A. Balbuena, Carl W. Dick, Katharina Dittmar, Susan Perkins

## Abstract

1. Ecological specialization reflects both evolutionary and ecological processes. For parasitic taxa, ecological specialization can be assessed as the degree to which a parasite species will associate with certain host species, a property known as host specificity.
2. Ectoparasitic bat flies have been previously reported as highly host specific, presumably due to a history of coevolution with their bat hosts. However, there is conflicting evidence of coevolution between bats and bat flies. Resource-driven competition between parasite individuals and between species may also be important in explaining patterns of bat fly specificity.
3. To test the importance of evolutionary and ecological factors on bat fly specificity, we collected and identified 21 bat fly species from 16 host bat species from the State of Rio de Janeiro in Brazil. We generated a bat fly species phylogeny from molecular data, estimated *d’* specialization values (a metric of specificity), and used linear and cophylogenetic models to compare the importance of various drivers of parasite ecological specialization.
4. We found that bat fly co-occurrence frequency (a proxy for interspecific competition) and mean infection intensity (a proxy for intraspecific competition) best predicted patterns of bat fly specialization. Co-occurrence frequency had a significantly negative association with specialization, while mean infection intensity has a significantly positive association with specialization. Coevolutionary congruence had a small effect size and did not significantly predict parasite specialization.
5. We found multiple shifts toward more generalized host niches across the bat fly phylogeny. Our results suggest that ecological processes such as resource-driven competition may be more important than evolutionary processes in shaping bat fly host specialization networks.
6. Bat flies showed variable degrees of host specialization, parasitized phylogenetically distant host species, and showed low phylogenetic congruence to their hosts. This suggests that as a group, bat flies may show flexibility in their host preference phenotypes and may change their host associations in the face of environmental disturbance.

## Introduction

Ecological specialization occurs when species preferentially occupy a niche as a result of natural selection, environmental adaptation, and/or interactions with other species (Blüthgen et al. 2006; Poulin et al. 2011; Clare et al. 2018). Ecological specialization has been demonstrated in plant-pollinator interactions (Blüthgen et al. 2006), feeding guilds occupied by animals (Rojas et al. 2011), gut microbiome communities (Ingala et al. 2018), and infection of hosts by parasites (Hatcher and Dunn 2011). Interactions between parasites and their hosts can serve as particularly good models for investigating ecological specialization because host-parasite interactions reflect the physical, ecological, and evolutionary relationships shared by both host and parasite. Parasites may be capable of infecting multiple host species, but often in unequal proportions and showing context-dependent variability (Marshall 1981; Rivera-García et al. 2017). Moreover, the suite of potential hosts utilized by a parasite species is governed by many factors including shared evolutionary history, morphological and immunological compatibility, ecologies of both parasite and host, competition between parasites for host resources, and the local environment (Marshall 1981; Hatcher and Dunn 2011; Poulin et al. 2011; Rivera-García et al. 2017; Zarazúa-Carbajal et al. 2016). Ultimately, the specificity of a parasite species to its host species may be determined by one or any combination of these factors. Studying host specialization in parasites provides a unique opportunity to examine how different ecological and evolutionary processes interact to produce patterns of host-parasite interactions and networks in biological communities.

Host specificity is a spectrum ranging from extremely host specific parasites (where a single parasite species occurs only on a single host species) at one end and generalist parasites (where a single parasite species occurs on multiple host species) at the other end. Extremely specific parasites tend to track the evolutionary history of their hosts due to a combination of coevolutionary mechanisms, life history traits such as reduced dispersal capability or obligate endoparasitism resulting from genetic and phenotypic divergence, and elevated selective pressures for the parasite to evolve highly specific adaptations to combat the host’s immune defenses (Reed and Hafner 2002; Poulin 2007). In contrast, many parasite species are less constrained and exhibit context-dependent associations with their hosts due to relaxed coevolutionary mechanisms and life history traits conducive to high levels of dispersal and mobility (e.g., some ectoparasites and parasite species with a multi-host life cycle have more opportunities for host-switching), resulting in evolutionary lability of host associations instead of specificity (Poulin et al. 2011; Galen et al. 2019). Because these different processes result in altered patterns of parasite specialization, we would expect highly specific parasites to show high phylogenetic congruence with their host species and, when those specific parasites do parasitize multiple hosts, to be restricted to phylogenetically related host species (Poulin et al. 2011; McQuaid and Britton 2013). Because generalist parasites may show evolutionary lability in their ability to parasitize multiple hosts, we would expect those parasites with reduced host specificity to show low phylogenetic congruence with their host species and to be capable of parasitizing many host species that are phylogenetically distant.

Evolutionary and ecological factors both contribute to the assembly of host-parasite communities (Lawton 1999; Poulin et al. 2011; Krasnov et al. 2011; Galen et al. 2019). For example, resource-driven competition between parasites, both at the intra- and interspecific levels, have been demonstrated in some parasitic taxa (Brown et al. 2009; Bush and Malenke 2008). In this circumstance, two parasite species utilizing the same host species can only coexist if they specialize to some degree on other host species, so ecological generalization would alleviate resource limitation in multi-host, multi-parasite systems (Dobson 1985). We would therefore expect parasite species that frequently share hosts with other parasite species to be generalists, and parasites that have excluded other species to be specialists (Dobson 1985; Dick and Dittmar 2014). Historical processes may also shape community composition based on whether traits associated with host preference are phylogenetically conserved or independently evolved (Cavender-Bares et al. 2004). If these traits are phylogenetically conserved, we would expect closely-related parasites to occur on the same host species or similar host species (Krasnov et al. 2016; Clark and Klegg 2017), while independently evolved host associations would promote communities of phylogenetically dissimilar parasites occupying a single host species (Galen et al. 2019). Historical and ecological processes may act in concert to influence host-parasite associations and therefore the ecological specialization of parasites.

To examine the interacting influences of ecology and evolution on host-associations, we used bat flies (Diptera: Streblidae, Nycteribiidae) as a model. Bat flies are obligate blood-feeding ectoparasites of bats that exhibit characteristics of both host specific and generalist parasites (Dick and Patterson 2007). Bat flies are extremely mobile and many species have functional wings, often moving between individuals of the same host species or leaving the host in order to ovoviviparously deposit larvae on roost substrate. However, bat flies must feed frequently and it is hypothesized that feeding preferences are extremely narrow (Dick 2007; Dick and Dittmar 2014). Previous studies of occurrence data indicate that bat flies have narrow, often 1-to-1 host associations (ter Hofstede et al. 2004; Dick and Patterson 2007; Dick 2007), but the single previous examination of bat and bat fly evolutionary associations indicates low phylogenetic congruence (Graciolli and de Carvalho 2012). Bat flies are uniquely suited to examine the impact of evolution and ecology on host specificity because they maintain traits necessary for both specialization and generalization, and bat fly species may be pushed towards one end of the host-preference spectrum by long-term associations, contemporary ecological pressures (e.g., competition), or environmental disturbance (Poulin et al. 2011; Pilosof et al. 2012).

Here, we examine the extent and drivers of host specialization of bat flies from the Atlantic Forest of Brazil. We use a combination of ecological networks, linear regression models, and cophylogenetic comparative methods to assess the following hypotheses:

1) If patterns of bat fly host specialization are the product of coevolutionary processes, cophylogenetic congruence with hosts should predict host specialization better than ecological predictors, because contemporary pressures of competition would be minimized in highly specialized parasites that have successfully excluded competing species. We should therefore expect instances of multi-host parasitism to be rare and, when they do occur, to be constrained to phylogenetically related hosts;
2) If bat flies instead show evolutionary lability in their host choice, we expect variables such as proxies of competition to better predict host specialization than cophylogenetic congruence. We should therefore expect bat fly species to be capable of parasitizing distantly related host species possibly due to phenotypic plasticity or convergent evolution of host preference in phylogenetically distant parasite species.

In this study, we examine ecological specialization of parasites at two distinct time scales; 1) the immediate ecological scale, in which observed host associations reflect recent host choice patterns, and 2) the deeper evolutionary scale, in which co-speciation events quantified through cophylogenetic methods indicate larger macroevolutionary processes that may contribute to extant patterns of host specialization of parasites. We argue that this combination of time scales is a useful strategy for examining the many factors that drive ecological specialization, beyond the scope of host-parasite interactions.

## Materials and Methods

### COLLECTION AND IDENTIFICATION OF PARASITES

From May through December, 2016, we sampled bats from 13 forest fragments within and around the Reserva Ecológica de Guapiaçu (REGUA), a mosaic of primary and secondary rainforest in the Atlantic Forest of Brazil (Rio de Janeiro State). REGUA encompasses a total protected area of 7,000 hectares. At its northern limit, it connects with the Serra dos Órgãos National Park and the Três Picos State Park to form one of the largest remnants of Atlantic Forest in Brazil (Supplementary Figure 1). The Atlantic Forest is one of the most heavily deforested areas in South America, with a long history of fragmentation (Bicudo da Silva et al. 2017).

For each fragment, we set up 7-10 mist nets for 6 hours every night for 6 nights. Sampling effort was similar for each fragment, ranging from 35-49 total nets set and 18,900-26,460 m^2^·h per fragment. We removed bats from mist nets and placed them in clean, individual cloth bags. After species identification of bats, we searched each individual for ectoparasites for a minimum of 45 seconds and collected as many observable ectoparasites as possible with forceps. We stored ectoparasites in 96% ethanol for transport and preservation. After each netting night, we washed all cloth bags to avoid parasite contamination into the next netting night.

We morphologically identified bat flies to the species level under a standard compound microscope using dichotomous keys specific to bat flies of the Neotropics (Wenzel 1976; Graciolli and Carvalho 2001a,b). For flies whose exoskeleton was damaged during collection or in transit, we assigned a tentative morphological identification, then reclassified the individual to species based on comparison with genetic clades after DNA sequencing (see below). After species identification, we constructed a database listing each individual bat fly with the corresponding host bat species from which it was collected (Supplement 1), and we calculated interaction frequencies from this database.

To confirm morphological identifications, we extracted genomic DNA from bat flies using DNeasy Tissue Extraction Kit (QIAGEN Inc) or Zymo Research ZymoBIOMICS DNA Miniprep Kit (Zymo Research, Irvine, CA). Prior to DNA extraction, each fly was separated into an individual microcentrifuge tube and digested in Proteinase K overnight. Extractions followed manufacturer protocol with some exceptions (see Supplement 2). We amplified a 645-basepair region of the cytochrome oxidase-I (*coxI*) mitochondrial gene using forward primer LC01490 and reverse primer HC02198 (Folmer et al. 1994) and a PCR protocol tailored from Hebert et al. (2003, 2004). We also amplified a 742-basepair region of the carbamoyl-phosphate synthetase (CAD) nuclear gene using a nested primer method with forward primer 787F and reverse primer 1124R based on methods described in Peterson et al. (2007). We visualized PCR products on 1.5% agarose gel through electrophoresis and cleaned PCR amplicons using AMPure XP beads (Beckman Coulter, Inc.). We cycle sequenced clean PCR products in 10μL reactions using Big Dye Terminator v3.1 (Life Technologies Corporation, Carlsbad, CA) chemistry following the standardly used protocol. Reactions were cleaned using AMPure XP beads prior to sequencing on a 3730xl Applied Biosystems (ABI) machine (Thermo Fisher Scientific). Primer and amplification details can be found in Supplement 2.

### DATA ANALYSIS

#### SPECIFICITY AND INTERACTION NETWORKS

From the database of individual parasite-host associations, we constructed a frequency matrix of the occurrence of parasite species on host species. This matrix was used to construct a bipartite interaction plot to visualize interaction frequencies using the R package *bipartite* (Dormann et al. 2008; R Core Team 2020).

Using the frequency matrix, we calculated the host specificity for each bat fly species, which is the total number of different host species with which a parasite is known to be associated within our dataset. The greater the number of host species utilized, the lower the specificity of that parasite species (Poulin et al. 2011). Although specificity is frequently reported in parasite studies, it is limited in that it does not incorporate frequency, abundance, phylogenetic distance, or sampling intensity for either hosts or parasites (Poulin et al. 2011). Specificity calculations also assume that parasites utilize all potential host species equally, which is often not the case in natural systems. Additionally, because highly specialized parasite species will all show a specificity value of 1.0, fine-scale statistical comparison between species that fall into this “super-specialists” group becomes impossible due to a lack of variability. To address these limitations, we calculated a frequency-based specialization metric, the *d’* index, for each parasite species, as well as its whole-network equivalent, the *H2* index (Blüthgen et al. 2006). The *d’* index attempts to correct for the assumption of uniform host preference by incorporating abundance data and frequency of host-parasite interactions. The *d’* index is scale invariant, so it allows for comparisons of parasite specificity across multiple spatial and time scales. The *d’* index produces a metric of host specificity ranging from 0 (perfect generalization) to 1 (perfect specialization) by quantifying the frequency distribution of the parasite’s niche utilization: a parasite species that uses all available host species proportionately to the availability of those hosts in the community would be considered more generalized in its niche utilization than a parasite species that utilizes rare hosts disproportionately (Blüthgen et al. 2006). The *d’* specialization index can therefore be used in multi-species parasite networks that differ in parasite species abundance due to sampling effort or rarity.

For each parasite species, we calculated the proportion of individual host infections that co-occurred with at least one other parasite species, a metric hereafter referred to as the co-occurrence frequency (COF). We used this as a proxy for the potential presence of interspecific competition, as parasite species with a higher COF are expected to co-occur with other parasite species more often than those with a lower COF. We also calculated the mean infection intensity (MI) for each parasite species, which is the mean number of individuals of that parasite species present per infected host individual (Aguiar and Antonini 2016). This MI metric serves as an estimate of the intraspecific densities that each parasite species encounters while parasitizing their respective hosts. For each parasite species, we calculated the phylogenetic distance between its host species using the “psv” command in R package *picante* (Kembel et al. 2010).

#### DRIVERS OF *D’* SPECIALIZATION

To test drivers of *d’* specialization, we constructed a global linear regression model including all predictor variables in the analysis: total fly abundance, number of hosts parasitized for each fly species (i.e., specificity), mean Rp_i_ (see below), COF, MI, and phylogenetic distance between host species. All covariates sufficiently met assumptions of normality after constructing histograms of residuals and conducting Kolmogorov-Smirnov tests in R (Massey 1951). We conducted multivariate tests for the presence of outliers and influential data points using the “outlierTest” and “cooks.distance” functions in the *car* package in R (Fox and Weisberg 2019). When all covariates were considered, we found one highly influential outlier data point, the bat fly species *Paraeuctenodes similis*, and it was subsequently excluded from the regression analyses. We generated all possible model combinations of predictor variables using the “dredge” function in the R package *MuMIn* (Bartoń 2019) and selected the top seven models based on their second-order Akaike Information Criterion (AICc, corrected for small sample size). The seven models chosen were then re-run, and the difference between each AICc, the minimum AICc value in the dataset (ΔAICc), and the importance weight for each model were calculated. Values of ΔAICc were considered significant when ≤ 2.0 (Burnham and Anderson 2004). We then partitioned and visualized variance explained by each covariate in the top selected model(s) using the “varpart” function in the R package *vegan* (Oksanen et al. 2019) and performed one-way ANOVAs for each covariate (compared to controlled effects of other covariates in model) to determine differences in effects on *d’* specialization. P-values for ANOVAs were calculated from 999 permutations. We visualized covariate effect sizes in forest plots using the “plot_summs” function in the R package *jtools* (Long 2019).

#### PHYLOGENETIC AND COEVOLUTIONARY ANALYSIS

For phylogenetic analyses, we first trimmed *coxI* and CAD sequences and manually corrected ambiguous nucleotide calls, generated consensus sequences for each parasite species, and aligned all sequences via ClustalW in GENEIOUS v. 10.1.2 (Biomatters Ltd.). We constructed a bat fly species tree with BEAST v. 2.5.1 and TreeAnnotator v. 2.5.1 using a gamma site model and GTR substitution model, a relaxed log normal clock model, a Yule pure speciation model prior, and a chain length of 50,000,000 generations and a 10,000-generation 2burn-in. Each gene was entered as a separate partition. The resulting species tree was used to quantify cophylogenetic congruence with the host bat phylogeny, which was trimmed from Shi and Rabosky’s (2015) larger phylogenetic tree of bats. We used the Random Tanglegram Partitions (Random TaPas) algorithm and Procrustean Approach to Cophylogeny (PACo, Balbuena et al. 2013; Balbuena et al. 2020), which construct Procrustes superimposition plots to test the phylogenetic congruence of parasite and host trees. The Random TaPas analysis yields a residual statistic (Rp_i_) for each host-parasite association that is proportional to cophylogenetic congruence. If the Rp_i_ is greater than zero, that host-parasite link is represented more often than expected by chance and is therefore phylogenetically congruent. If the interaction Rp_i_ is less than zero, that host-parasite link is represented less often than expected by chance and is therefore phylogenetically incongruent. The Random TaPas analysis also estimates ancestral states of each node using fast maximum likelihood estimation, elucidating specialization switching patterns across the parasite phylogeny. We calculated the Rp_i_ for each host-parasite combination in our dataset using our bat fly phylogeny and trimmed bat phylogeny. This Rp_i_ was then mapped onto a host-parasite tanglegram to illustrate the strength of phylogenetic congruence across the host-parasite cophylogeny. We quantified transitions from ecological generalization to specialization (and vice-versa) for terminal parasite taxa by comparing estimated ancestral states; clade transitions in which the derived node showed higher host congruence than the ancestral node was considered to be a transition to host specialization, whereas instances in which the derived node showed lower congruence than the ancestral node was considered a transition to ecological generalization.

## Results

### SPECIALIZATION AND INTERACTION NETWORKS

We identified 21 bat fly species (832 individuals) from 16 bat host species (360 individuals, Table 1, Fig 1). Specificity values ranged from one to five bat host species per parasite species with a mean of 1.9 and a standard deviation (sd) of 1.2 (Table 1). The *d’* specialization indices for our bat fly species ranged from 0.080 (*Paraeuctenodes similis*) to 1.0 (*Basilia juquiensis, Neotrichobius delicatus, Trichobius longipes, Trichobius* sp. 2) with a mean value of 0.713 (sd = 0.31). The *H2* index for entire community specialization was 0.889. For those parasites that were found associated with more than one host species, the mean phylogenetic distance between host species was 0.40 (sd = 0.16), indicating that bat flies with a wider niche breadth are not restricted to closely phylogenetically related host species. In fact, only three (*Aspidoptera falcata, Aspidoptera phyllostomatis,* and *Basilia juquiensis*) of the ten bat fly species that parasitized multiple host species were restricted to congeneric hosts. The remaining seven fly species parasitized hosts of non-sister clades (Figs 1, 2).

**Figure 1.**
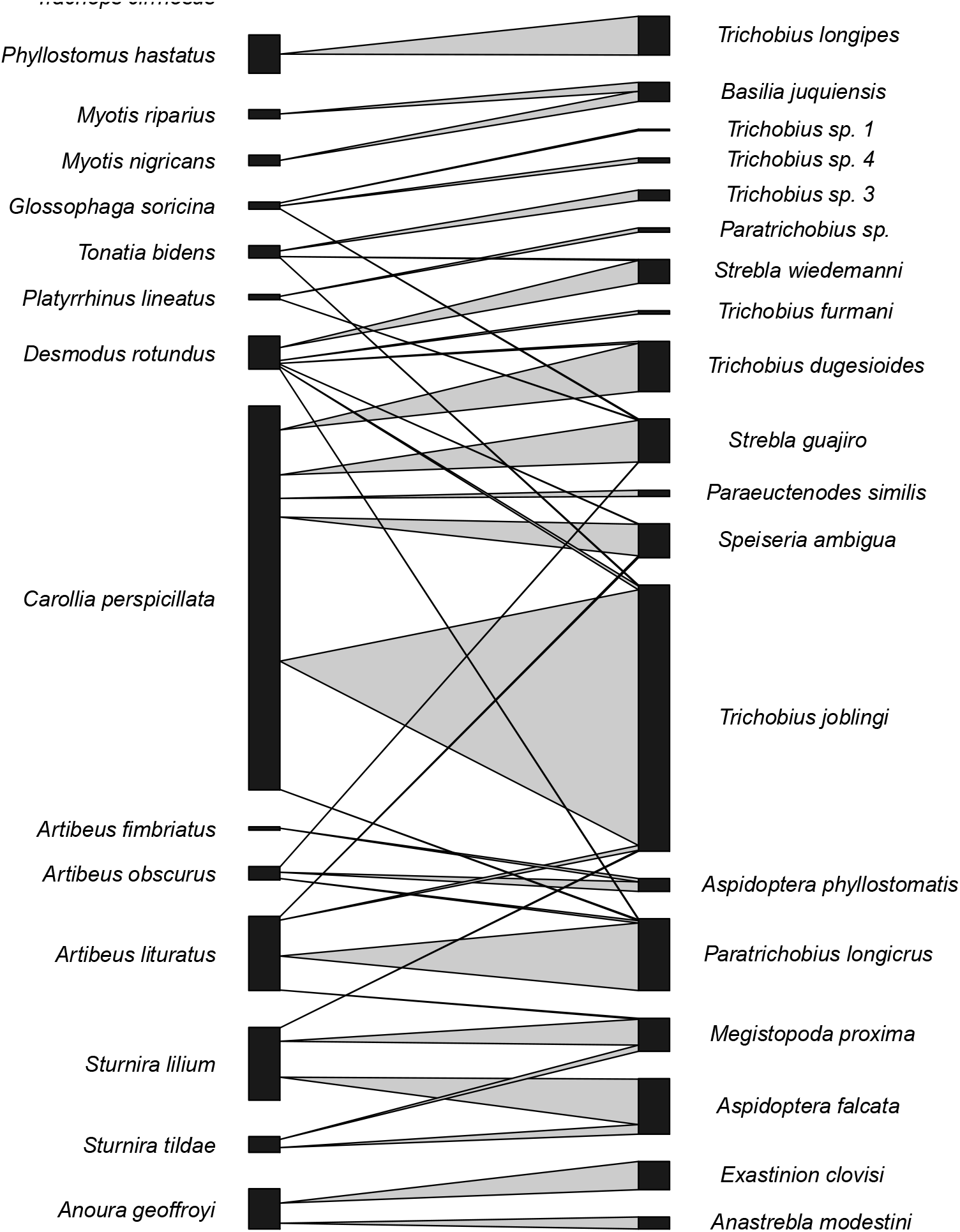
Bipartite interaction network for all bat host species (left) and all bat fly parasite species (right). Lines connecting parasites and hosts represent infection of that host by that parasite, with the thickness reflecting the frequency of the interaction. Heights of species boxes reflect sample abundances.

**Table 1.** Total abundance (N), *d’* specialization (*d’*), host specificity (B), host genetic distance (P), co-occurrence frequency (COF), and mean infection intensity (MI) values for each of the bat fly species identified.

| Bat Fly Species | N | $d'$ | B | P | COF | MI |
| --- | --- | --- | --- | --- | --- | --- |
| *Anastrebla modestini* | 14 | 0.698 | 1 | 0.000 | 0.428 | 2.000 |
| *Aspidoptera falcata* | 63 | 0.816 | 2 | 0.153 | 0.542 | 2.750 |
| *Aspidoptera phyllostomatis* | 15 | 0.941 | 2 | 0.142 | 0.000 | 2.140 |
| *Basilia juquiensis* | 22 | 1.000 | 2 | 0.297 | 0.000 | 2.750 |
| *Exastinion clovisi* | 32 | 0.886 | 1 | 0.000 | 0.714 | 4.570 |
| *Megistopoda proxima* | 38 | 0.639 | 2 | 0.319 | 0.482 | 1.410 |
| *Neotrichobius delicatus* | 6 | 1.000 | 1 | 0.00 | 0.000 | 1.500 |
| *Paraeuctenodes similis* | 7 | 0.080 | 1 | 0.000 | 0.167 | 1.170 |
| *Paratrichobius longicrus* | 82 | 0.883 | 4 | 0.454 | 0.000 | 1.577 |
| *Paratrichobius* sp | 5 | 0.962 | 1 | 0.000 | 0.000 | 2.500 |
| *Speiseria ambigua* | 39 | 0.139 | 4 | 0.531 | 0.778 | 1.220 |
| *Strebla guajiro* | 50 | 0.197 | 5 | 0.469 | 0.641 | 1.250 |
| *Strebla wiedemanni* | 27 | 0.861 | 2 | 0.563 | 0.167 | 1.420 |
| *Trichobius dugesioides* | 57 | 0.206 | 4 | 0.563 | 0.727 | 1.270 |
| *Trichobius furmani* | 4 | 0.541 | 1 | 0.000 | 0.667 | 1.330 |
| *Trichobius joblingi* | 303 | 0.514 | 5 | 0.511 | 0.386 | 2.180 |
| *Trichobius longipes* | 44 | 1.000 | 1 | 0.000 | 0.000 | 6.290 |
| *Trichobius* sp 1 | 2 | 0.743 | 1 | 0.000 | 0.000 | 2.000 |
| *Trichobius* sp 2 | 5 | 1.000 | 1 | 0.000 | 0.000 | 5.000 |
| *Trichobius* sp 3 | 12 | 0.962 | 1 | 0.000 | 1.000 | 12.00 |
| *Trichobius* sp 4 | 5 | 0.901 | 1 | 0.000 | 0.500 | 1.250 |

From the Random TaPas analysis, we determined 40 unique bat fly-bat host associations (Table 2). Interaction frequencies ranged from 1 to 298 and Rp_i_ ranged from -28.49 (low congruence between the bat *Artibeus fimbriatus* and bat fly *Aspidoptera phyllostomatis*) to 34.72 (high congruence: *Anoura geoffroyi – Exastinion clovisi*) (Table 2). The mean Rp_i_ value for all interactions was 0.10 (standard deviation = 13.3), indicating that the majority of interactions showed low levels of phylogenetic congruence between host and parasite (Fig 2). From the ancestral state estimations, we quantified 6 total transitions from host specialization to generalization, and 7 transitions from host generalization to specialization for terminal parasite taxa.

**Table 2.**
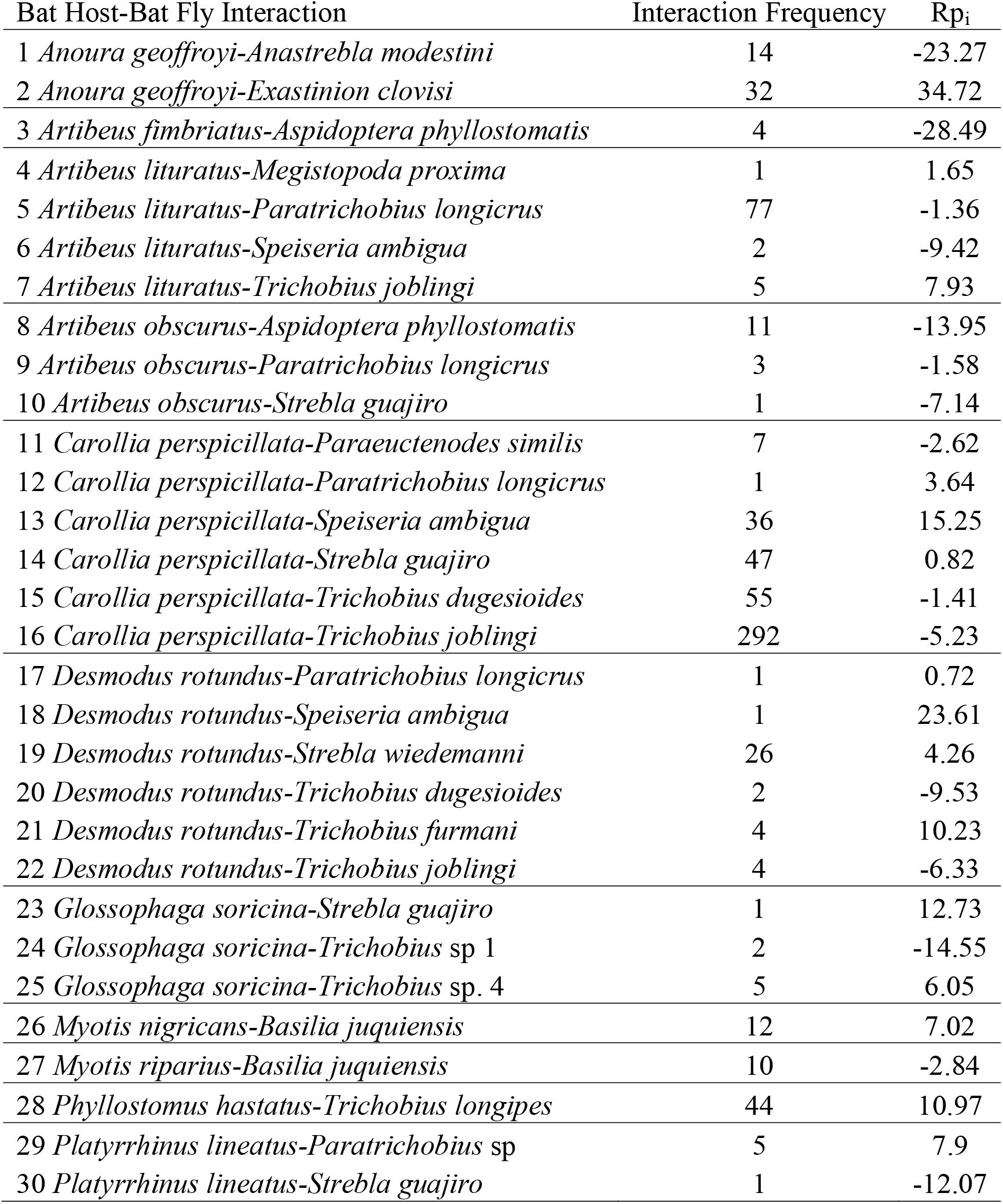
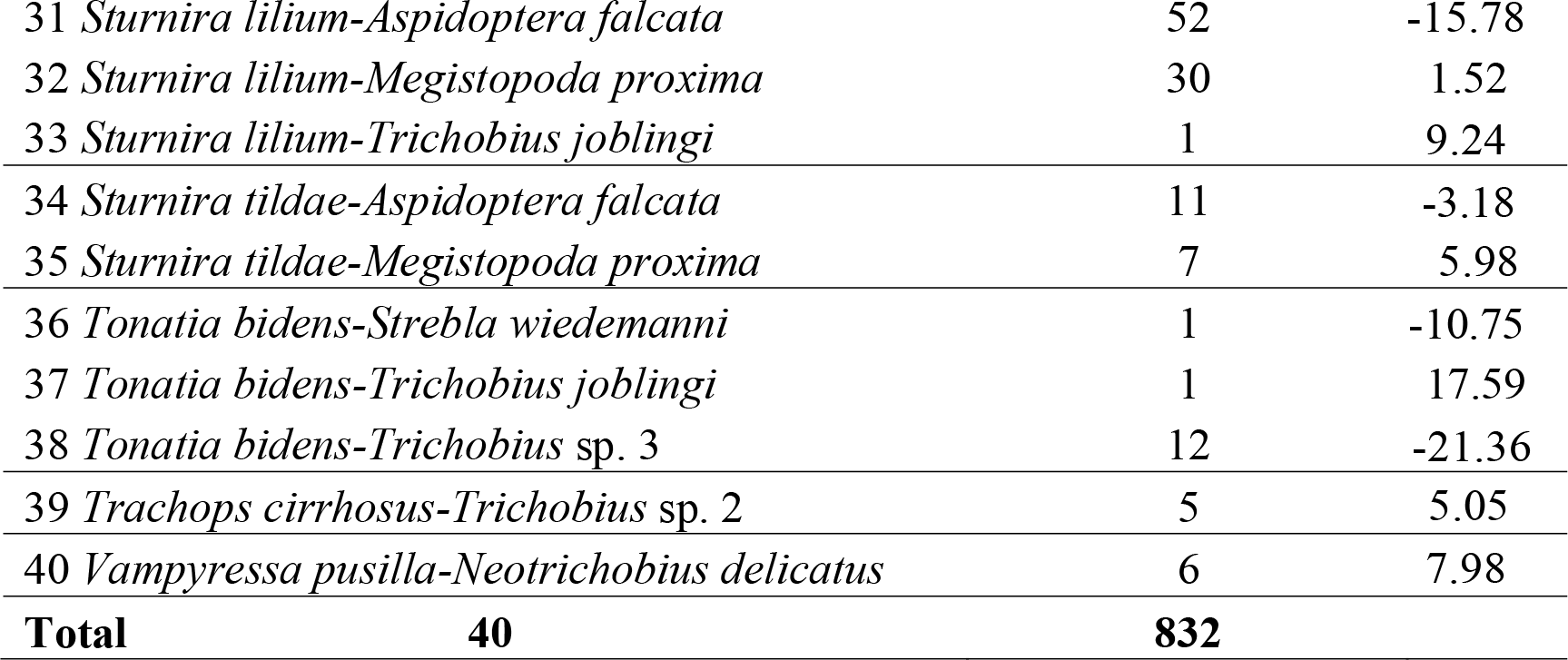
Interaction frequency and Random TaPas Rp_i_ values for each bat fly-bat host interaction observed in the whole-community bipartite network. Rp_i_ values are directly proportionate to phylogenetic congruence; higher values reflect a greater parasite phylogeny congruence with its host’s phylogeny.

**Figure 2.**
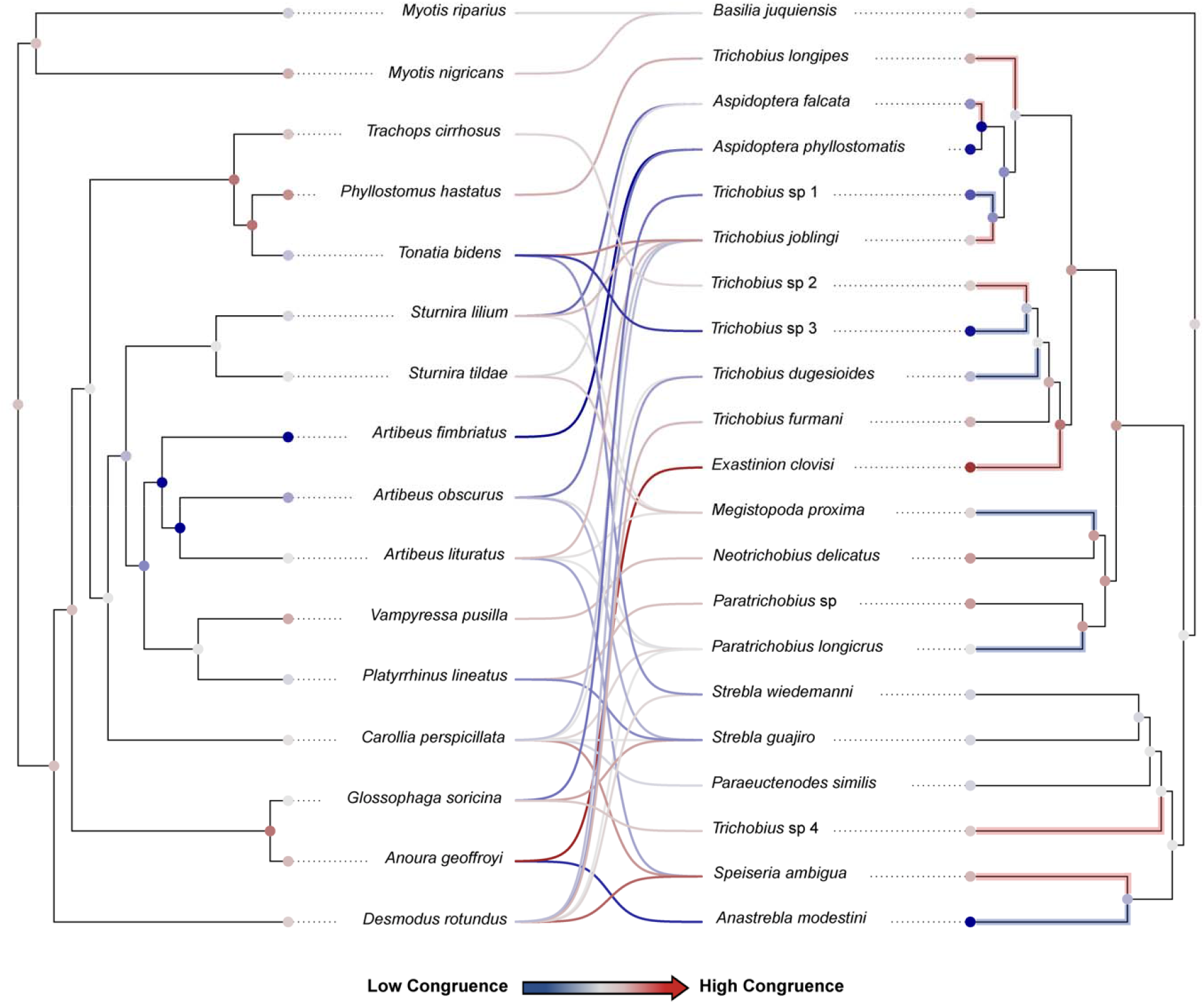
Tanglegram of bat host species (left) and their associated bat fly species (right) generated from the Random TaPas analysis. Connecting lines represent an interaction, and colors of lines represent mean Rp_i_, or strength of congruence between host and parasite tips. Color scheme ranges from dark blue (weak congruence) to dark red (strong congruence). Parasite branches for which a shift to a more generalized (blue) or a more specialized (red) host niche from the ancestral state are highlighted. Node colors reflect ancestral state estimation based on rapid ML estimation methods.

### DRIVERS OF *d’* SPECIALIZATION

From the global model selection analysis, *d’* specialization was best predicted by model *d’* ∼ COF + MI + P (AICc = -7.74, ΔAICc = 0.0, weight = 0.45, Table 3) and model *d’* ∼ COF + MI + B (AICc = -6.17, ΔAICc = 1.58, weight = 0.20, Table 3). Mean effect estimates for COF, P, and B were all negative; *d’* specialization increased as COF, P, and B decreased, and vice-versa. In contrast, effect sizes for MI were positive; *d’* specialization decreased and increased as MI decreased and increased (Fig 3). Effect size for total fly abundance (A) was negative and effect size for mean Rp_i_ was positive, but with lower confidence (95% CIs for estimates included zero).

**Figure 3.**
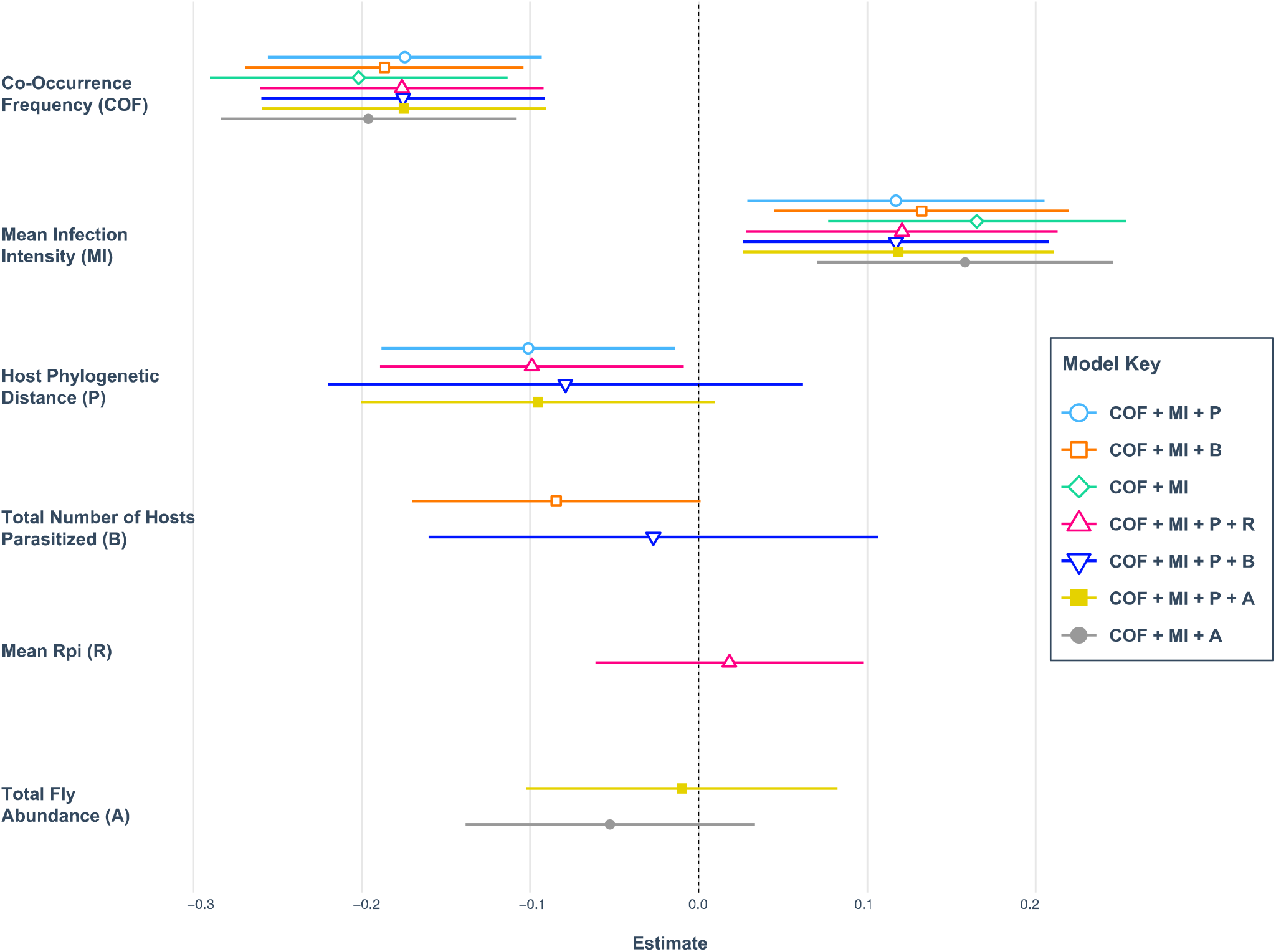
Forest plot depicting regression outputs for the best 7 models predicting *d’* specialization for all bat fly species. Models are differentiated by bar colors and shape of the mean estimate point nested within 95% confidence interval bars. Predictor variables are listed on the y-axis, with colored bars adjacent to each predictor variable representing regression models in which that predictor is included. The vertical dashed line indicates the zero-effect estimate. Estimates falling to the left of the dashed line indicate a negative effect of that predictor variable on *d’* specialization; bars falling to the right indicate a positive effect.

**Table 3.** Model comparison statistics from linear regressions performed on *d’* specialization for the whole bat fly community including all fly species. Model formula term abbreviations are as follows: A = total parasite abundance, B = total number of hosts parasitized (specificity), COF = co-occurrence frequency, MI = mean infection intensity per parasite species, R= mean Rp_i_, P = phylogenetic distance between host bat species.

| Model Predictors | df | Deviance | AICc | ΔAICc | Weight | $R^2$ |
| --- | --- | --- | --- | --- | --- | --- |
| COF + MI + P | 5 | 0.39 | -7.74 | 0.00 | 0.45 | 0.75 |
| COF + MI + B | 5 | 0.42 | -6.17 | 1.58 | 0.20 | 0.73 |
| COF + MI | 4 | 0.54 | -4.94 | 2.80 | 0.11 | 0.65 |
| COF + MI + P + R | 6 | 0.38 | -3.88 | 3.86 | 0.07 | 0.75 |
| COF + MI + B + P | 6 | 0.38 | -3.81 | 3.93 | 0.06 | 0.75 |
| COF + MI + P + A | 6 | 0.39 | -3.64 | 4.11 | 0.06 | 0.75 |
| COF + MI + A | 5 | 0.49 | -3.34 | 4.40 | 0.05 | 0.69 |

Total variance in *d’* specialization explained by covariates COI, MI, P, and B included in the top two regression models above (adjusted R^2^) was 0.684 (residual variation = 0.316. Fig 4). After variance partitioning of covariates, COF accounted for the greatest proportion of explanatory variance (R^2^ = 0.3705), followed by MI (R^2^ = 0.1285), P (R^2^ = 0.0085), and B (R^2^ = – 0.016; Fig 4). After performing one-way ANOVAs to test for differences in effects of top model covariates, only COI (p = 0.002) and MI (p = 0.012) were statistically significant (Table 4).

**Figure 4.**
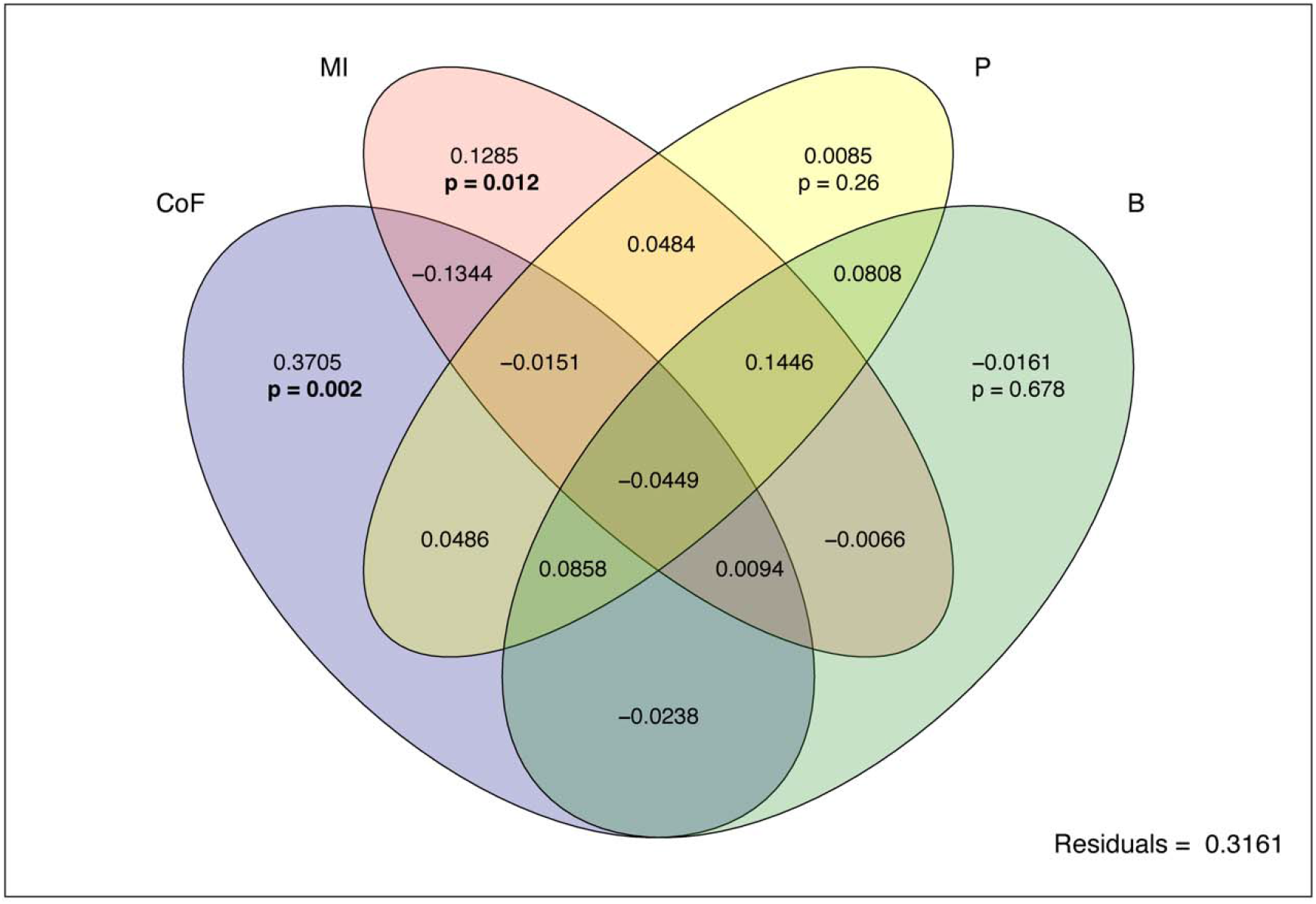
Venn diagram depicting total variance in *d’* specialization explained by covariates in the top two AICc-selected regression models, partitioned by the partial R^2^ value for each covariate. Note that because partial R^2^ for each covariate was calculated by controlling for the other three, some partition values may be negative and are considered uninterpretable. P-values from ANOVA were calculated with 999 permutations.

**Table 4.** One-way ANOVA table for differences in effect of covariates included in the top two models explaining *d’* specialization. P-values were calculated from 999 permutations.

| Covariate | df | F-statistic | p-value |
| --- | --- | --- | --- |
| COI | 1 | 19.75 | **0.002\*\*** |
| MI | 1 | 7.51 | **0.012\*** |
| P | 1 | 1.43 | 0.26 |
| B | 1 | 0.185 | 0.678 |

## Discussion

In this study, we hypothesized that if extant host associations of bat flies have evolved due to cospeciation, then variation in *d’* specialization of bat flies should be best explained by mean Rp_i_ (phylogenetic congruence) and multi-host parasitism should be constrained to phylogenetically related hosts. Conversely, if bat fly host choice is determined by contemporary ecological pressures such as competition, then ecological variables such as COF and MI should best explain variation in *d’* specialization and bat flies should parasitize phylogenetically distant hosts. Our results demonstrate that bat flies are not highly specialized as a group — generalist fly species can and do parasitize phylogenetically distant host species, and shared evolutionary history with host bats is not the main force shaping bat fly niche breadth. Instead, host specialization was best explained by co-occurrence frequency and mean infection intensity, suggesting that fine-scale interspecific interactions between bat fly species may be more important in dictating extant patterns of bat fly host specialization than evolutionary processes. Moreover, we demonstrate that host specialization and generalization appear to have evolved independently across multiple lineages, indicating that traits associated with host preference are not phylogenetically conserved in bat fly clades. Ours is the first study to test host specialization patterns of bat flies at both the deep evolutionary scale and the contemporary ecological scale, and has important implications for studying how host-parasite networks can be affected by a variety of interrelated processes. This work contributes to the growing body of research illustrating that both evolutionary and ecological processes impact host-parasite interactions (Choisy and de Roode 2010; Galen et al. 2019).

Many of the fly species in our study showed fairly low *d’* specialization values, indicating that not all bat fly species preferentially specialize on a singular host species. Although this finding seemingly contradicts previous studies (ter Hofstede et al. 2004; Dick 2007; Dick and Patterson 2007), a trend towards ecological generalization in bat flies is consistent with bat fly biology; bat flies are mobile ectoparasites that are capable of flying from one host individual to another and/or from the host individual to the roost substrate, particularly in the case of reproductive females preparing to deposit pupae (Dittmar et al. 2009). Ecological theory predicts that a generalist host niche should be favored by a parasite if the benefits of parasitizing multiple host species (increased potential host density, higher probability of encountering mates, reduced probability of local extinction) outweigh the costs of forgoing specialization (for example, increased performance resulting from adaptive specialization to a single host) (Dobson 1985; Futuyma and Moreno 1988; Agosta et al. 2010). Because bat flies are ectoparasites, they live on the host’s external surfaces and are thus not immediately subject to the host’s innate immune response. This may reduce selective pressure to evolve specialized adaptations to combat host immune defenses, potentially resulting in a trend toward more generalized life history strategies including wider niche breadth (Poulin et al. 2011; Pinheiro et al. 2016). Bat flies have short lifespans and will typically die within 24 hours if removed from their host (Dick and Dittmar 2014). Additionally, most bat flies will reach sexual maturity 5-6 days after puparium emergence and will immediately search for a mate (Dittmar et al. 2009). It is possible that fitness pressures to consistently obtain a blood meal and quickly find a mate could greatly outweigh the potential value of specializing on a single host, resulting in ecologically generalized feeding niches (Poulin 2007; Dick and Dittmar 2014; Pinheiro et al. 2016).

It is important to note that we did encounter uncertainties regarding the validity of some of our observed host-parasite associations. For example, of our 40 host-parasite interactions, 14 of these indicate a parasite associated with a “nonprimary” host as defined by Dick and Patterson (2007). However, 10 of these 14 “nonprimary” associations have been reported in other scientific studies of bat flies in Brazil (Graciolli 2001; Camilotti et al. 2010; Barbier and Graciolli 2016; de Vasconcelos et al. 2016; Aguiar and Antonini 2016), supporting their validity. By using a specialization metric which includes information about frequency of the association, we limit the impact of potential contamination in our data in driving measures of specificity. When we removed the 14 “nonprimary” associations from our dataset and recalculated *d’* specialization values for all parasite species, there was no significant difference in specialization values from the original dataset that included the “nonprimary” associations (t = 0, df = 41.99, p = 1). This lack of statistical difference, in conjunction with support from other studies of bat fly specificity from our specific study region, encouraged us to include our “nonprimary” associations in our analysis rather than exclude them, because a conservative approach that excludes certain host-parasite associations may risk under-sampling the range of bat fly host specialization (Poulin et al. 2011).

The ecology of host bats themselves is also likely to influence a trend towards generalized feeding niches in bat flies. Tropical bat species are unique in their abundances and roost-switching behaviors, often migrating over large distances in short periods of time and sometimes occupying different roosts on a nightly basis (Graham 1988; Kunz and Fenton 2005; Voss et al. 2016). Frequent roost switching by bats can result in high rates of turnover of bat species assemblages available to bat fly pupae developing on roost substrate. If bat flies in this scenario were highly specialized to a single host species, it would increase their chance of local extinction if that particular host left the roost and was not replaced by new individuals of the same species. It would incur a greater fitness advantage for bat flies to show phenotypic plasticity in their ability to parasitize a wide range of host species, given the somewhat unpredictable nature of roost switching by host bats (Graham 1988; Voss et al. 2016). We found an interesting association in our study that may be evidence of this phenomenon; *Strebla wiedemanni* commonly parasitizes *Desmodus rotundus* (Wenzel 1975), but we also observed *Strebla wiedemanni* parasitizing *Tonatia bidens*. *Tonatia bidens* and *Desmodus rotundus* are bat species that have similar roosting ecologies (Voss et al. 2016) and could potentially share fly species if they are sharing roosts in our study area. In addition, *Strebla wiedemanni* showed a relatively weak coevolutionary signal with its primary host species, *Desmodus rotundus* (Rp_i_ = 4.26), suggesting this bat fly species may show phenotypic plasticity in its host choice and could opportunistically parasitize atypical host species (such as *Tonatia bidens*) if *Desmodus rotundus* were not present.

Our cophylogenetic analysis further supported our interpretation that bat flies may be opportunistic generalists in their host choice. We found that the majority of interactions had low phylogenetic congruence between host and parasite taxa. This is consistent with the only other cophylogenetic study of bat flies and their hosts (Graciolli and de Carvalho 2012), but contradicts studies on host specificity that suggest bat flies evolve in tight association with their host bats (ter Hofstede et al. 2004; Dick 2007). The consistency of our findings with those of Graciolli and de Carvalho (2012) is interesting to note, as these authors investigated cophylogenetics of the genus *Trichobius* based only on morphological characters. The fact that bat flies show low cophylogenetic signal with their host bats may be evidence of evolutionary lability of host choice as a strategy to maximize fitness by broadening niche breadth (Agosta et al. 2010; Pinheiro et al. 2016). This hypothesis is consistent with our findings regarding the phylogenetic distance between hosts; of the 10 bat fly species which were associated with more than one host, only 3 of these species were restricted to congeneric hosts and the remaining 7 parasitized more phylogenetically distant hosts. Moreover, ancestral state estimations showed several independent host-switching events across multiple lineages, with 6 total extant taxa evolving more generalized host niche breadths than their ancestral state. If bat flies are not evolutionarily constrained in their host choice, they are likely to occupy a “sloppy fitness space” in which multiple hosts can be parasitized due to generalized utility of the parasite’s traits (Agosta et al. 2010). In this way, bat flies may actually be more specialized to particular resource (bat blood itself) than to the specific identity of the bat hosts, and frequent patterns of host switching and ecological generalization should be observed (Poulin 2007; Agosta et al. 2010; Pinheiro et al. 2016). Our results demonstrate a combination of low phylogenetic congruence, phylogenetically diverse host suites, and multiple host-switching events across bat fly lineages. These findings support our hypothesis that cospeciation is not the main process determining patterns of extant bat fly-bat host associations in tropical communities, and that bat flies likely show evolutionary lability in their host specialization niches.

The Atlantic Forest of Brazil is highly fragmented with a long history of deforestation and disturbance, which may explain trends towards ecological generalization in bat flies in the region. Because most of the bat species in our study roost in forest structures such as tree hollows (Graham 1988; Voss et al. 2016), anthropogenic modification of the forest landscape can affect roost availability and, by extension, local abundances and species compositions of bats (Medellín et al. 2000; Pilosof et al. 2012; Hiller et al. 2020). Disruptions in the bat community will undoubtedly affect host-parasite networks; if bat flies cannot parasitize a preferred host due to changes in bat abundances or species composition, bat flies will be forced either to parasitize other available host species or, if they are highly specialized and evolutionarily constrained in their host choice, go locally extinct (Dobson 1985; Futuyma and Moreno 1988; Pilosof et al. 2012). Most bat fly species in our study showed low phylogenetic congruence with their host species, suggesting that they are not evolutionarily constrained to a particular host suite and may opportunistically parasitize other hosts in the face of ecological disturbance. Additionally, host-switching was demonstrated to occur multiple times across the parasite phylogeny, and even sister parasite taxa showed divergent host associations from their ancestral state. This can be interpreted as evidence that bat flies may have the evolutionary lability to switch hosts and avoid local extinction should local bat assemblages be affected by anthropogenic disturbance (Vázquez and Simberloff 2002; Agosta et al. 2010).

Although we did not explicitly test this, bat fly species that tended to co-occur together on the same host were not closely related phylogenetically. For example, three bat fly species frequently co-occurred on *Carollia perspicillata: Trichobius joblingi, Strebla guajiro, and Speiseria ambigua*. These parasite species are in different genera, fall in completely separate clades in our phylogeny (see Fig 2), and are morphologically distinct (Wenzel 1976, Tello et al. 2008). Although they may co-occur on the same bat individual, this does not mean that they directly compete for resources. Species in the genus *Trichobius* tends to be found most frequently on bat wing membranes and interfemoral regions, while *Strebla* and *Speiseria* are known to occupy furred areas of the bat host’s body, an apparent strategy to reduce on-host competition for space and resources (ter Hofstede et al. 2004). This strategy may explain the patterns observed in our study and suggest bat flies may be capable of avoiding the effects of exploitation-mediated competition with other co-occurring bat fly species while also maintaining a broader ecological niche in terms of their suite of potential hosts (*d’* specialization). In this way, bat flies may be utilizing both broad-scale host generalization and small-scale site specialization to minimize the adverse effects of specialization tradeoffs (Dobson 1985; Futuyma and Moreno 1988; ter Hofstede et al. 2004). While co-occurrence frequency can serve as an indicator of the *potential* for interspecific competition between parasites, it cannot definitively indicate the presence or specific nature of this competition (Blanchet et al. 2020). To determine the extent of competition between bat fly species that may be driving ecological specialization, future research should include variation in parasite traits such as phylogenetic relatedness and morphological traits, as well as perform explicit null-model competition analyses (Tello et al. 2008, Blanchet et al. 2020).

In contrast to COF, mean infection intensity (MI) showed the opposite effect on *d’* specialization: bat fly species with higher *d’* specialization values tended to show higher on-host densities. This finding is consistent with predictions of specialization: as a parasite species becomes more ecologically specialized to one host species, its overall niche breadth (and potential to switch host species) is reduced and on-host intraspecific densities should subsequently increase (Dobson 1985; Hatcher and Dunn 2011). This observation may be evidence that specialized bat flies are competitively excluding other species on-host: many fly species in our study that showed high *d’* specialization values also showed low COF values (see Table 1). Additionally, some of these species that show low COF values have morphologies that may aid in competitively excluding other competing bat fly species. *Trichobius longipes,* a specialist parasite of *Phyllostomus hastatus,* is by far the largest bat fly species in our sample (body size can get up to 0.82 mm; Wenzel 1975), occurred in high intraspecific densities, and never occurred with another fly species. The size dominance of *Trichobius longipes* may be a contributing factor in its specialization to a single host species, given that larger parasites in high densities are likely to outcompete other smaller parasites for space and resources on the host’s body surface (Wenzel 1976; Dobson 1985; Hatcher and Dunn 2011; Presley 2011). This was not explicitly tested in our study but presents an interesting line of research for future studies.

## Conclusions

Our study has demonstrated that patterns of Neotropical bat fly parasitism are dependent on a variety of interrelated factors spanning the fields of ecological and evolutionary biology. Bat flies are a complex group of parasites that show a range of ecological specialization to their host bats. Ours is the first empirical study to quantitatively investigate patterns of bat fly host specialization in both a phylogenetic and ecological context. We provide evidence that both host specialization and cophylogenetic congruence are differentially distributed across the Streblidae. Bat fly species are apparently not constrained to tight cophylogenetic relationships with their hosts and may exhibit flexibility in their host choice, allowing for niche broadening when faced with ecological or environmental disturbance. Ecologically generalized fly species form multi-species assemblages on host individuals more frequently than specialized species, suggesting that resource partitioning may be driving host-parasite assemblages at the individual level, but not at the host specialization level. Specialist fly species show elevated levels of on-host population densities, possibly as a result of specialization reducing the suite of potential hosts that would allow for alleviated intraspecific competition.

The results of our study may have important implications for vector disease biology. By studying the multiple drivers of parasite specificity, and how they vary based on host and parasite evolutionary history, researchers may gain important insights useful for improving modeling of disease transmission between arthropods, wildlife, and potentially humans. Understanding the complex interactions among parasites and pathogen hosts and vectors is becoming increasingly important with the ongoing discovery of new wildlife diseases and continuing deforestation in areas like the Atlantic Forest, which is effectively minimizing natural buffer areas between wildlife disease vectors and human populations. Particularly given recent concerns about bats as potential reservoirs of zoonotic diseases (Brook et al. 2020; Ge et al. 2020; Zhou et al. 2020), understanding the ecology and evolution of blood-feeding bat ectoparasites is of considerable societal importance. Combining analytical methods that examine ecological specialization at multiple time scales (both ecological and evolutionary) has the potential to elucidate patterns that may not be immediately apparent with traditional metrics. These methods could be applied to a variety of biological disciplines other than parasitology, including pollination biology, microbiome evolution, and ecosystem ecology.

## Acknowledgements

We would like to thank our field and laboratory colleagues from Columbia University, the American Museum of Natural History, and Queen Mary University of London for contributing to this highly collaborative research. AMB received financial support from the American Museum of Natural History and Sackler Institute for Comparative Genomics to complete this research. KAS was funded through the Richard Gilder Graduate School Student Research Fellowship. Special thanks to JJB for analytical suggestions and moral support.

## Authors’ Contributions

AMB designed the study focus, identified parasites, conducted molecular work, performed analyses, and wrote the paper. KAS identified parasites, performed analyses, aided in molecular work, and contributed to writing the paper. TT performed field work, collected samples, and contributed field site coordinates and data information. SP provided laboratory reagents, aided in molecular work, and contributed to writing the paper. CWD and KD aided in parasite identification and contributed to writing the paper. EC, NBS, and JAB contributed to writing the paper.

## Data Availability Statement

All genetic data was deposited into NCBI GenBank (accession numbers forthcoming). All supplements, datasets, and code used in analyses will be available on Dryad Digital Repository (https://doi.org/10.5061/dryad.b2rbnzsdd) upon receipt of a manuscript DOI.

## References

Agosta, S. J., Janz, N., & Brooks, D. R. (2010). How specialists can be generalists: resolving the “parasite paradox” and implications for emerging infectious disease. Zoologia (Curitiba), 27(2), 151–162. https://doi.org/10.1590/S1984-46702010000200001

Aguiar, L., & Antonini, Y. (2016). Prevalence and intensity of Streblidae in bats from a Neotropical savanna region in Brazil. Folia Parasitologica, 63(1). doi: 10.14411/fp.2016.024

Balbuena, J., Míguez-Lozano, R., & Blasco-Costa, I. (2013). PACo: A Novel Procrustes Application to Cophylogenetic Analysis. PloS One, 8(4): e61048. doi: 10.1371/journal.pone.0061048

Balbuena, J. A., Pérez-Escobar, Ó. A., Llopis-Belenguer, C., & Blasco-Costa, I. (2020). Random Tanglegram Partitions (Random TaPas): An Alexandrian Approach to the Cophylogenetic Gordian Knot. Systematic Biology, syaa 033. doi: 10.1093/sysbio/syaa033 (In press).

Barbier E., & Graciolli, G. (2016). Community of bat flies (Streblidae and Nycteribiidae) on bats in the Cerrado of Central-West Brazil: hosts, aggregation, prevalence, infestation intensity, and infracommunities. Studies on Neotropical Fauna and Environment, 51(3): 176–187. doi: http://dx.doi.org.ezproxy.cul.columbia.edu/ 10.1080/01650521.2016.1215042

Barto, K. (2019). MuMIn: Multi-Model Inference. R package version 1.43.6. ń https://CRAN.R-project.org/package=MuMIn

Bicudo da Silva, R. F., Batistella, M., Moran, E. F., & Lu, D. (2017). Land Changes Fostering Atlantic Forest Transition in Brazil: Evidence from the Paraíba Valley. The Professional Geographer, 69(1), 80–93. doi: 10.1080/00330124.2016.1178151

Blanchet, F. G., Cazelles, K., & Gravel, D. (2020). Co-occurrence is not evidence of ecological interactions. Ecology Letters, 23(7), 1050–1063. doi: 10.1111/ele.13525

Blüthgen, N., Menzel, F., & Blüthgen, N. (2006). Measuring specialization in species interaction networks. BMC Ecology, 6(1), 9. doi: 10.1186/1472-6785-6-9

Brook, C. E., Boots, M., Chandran, K., Dobson, A. P., Drosten, C., Graham, A. L., Grenfell, B. T., Müller, M. A., Ng, M., Wang, L.-F., & van Leeuwen, A. (2020). Accelerated viral dynamics in bat cell lines, with implications for zoonotic emergence. ELife, 9, e48401. https://doi.org/10.7554/eLife.48401

Brown, S. P., Inglis, R. F., & Taddei, F. (2009). Evolutionary ecology of microbial wars: within-host competition and (incidental) virulence. Evolutionary Applications, 2(1): 32–39. doi: 10.1111/j.1752-4571.2008.00059.x

Burnham, K., & Anderson, D. (2004). Multimodel Inference: Understanding AIC and BIC in Model Selection. Sociological Methods & Research, 33(2): 261–304. doi: 10.1177/0049124104268644

Bush, S. E., & Malenke, J. R. (2008). Host defence mediates interspecific competition in ectoparasites. Journal of Animal Ecology, 77(3): 558–564. doi: 10.1111/j.1365-2656.2007.01353.x

Camilotti, V., Graciolli, G., Weber, M., Arruda, J., & Cáceres, N. (2010). Bat flies from the deciduous Atlantic Forest in southern Brazil: Host-parasite relationships and parasitism rates. Acta Parasitologica, 55(2): 194–200. doi: 10.2478/s11686-010-0026-2

Cavender-Bares, J., Ackerly, D.D., Baum, D.A., & Bazzaz, F.A. (2004). Phylogenetic overdispersion in Floridian oak. The American Naturalist, 163(6), 823–843. https://doi.org/10.1086/386375

Choisy, M., & de Roode, J. C. (2010). Mixed Infections and the Evolution of Virulence: Effects of Resource Competition, Parasite Plasticity, and Impaired Host Immunity. The American Naturalist, 175(5), E105–E118. https://doi.org/10.1086/651587

Clare, E. L., Fazekas, A. J., Ivanova, N. V., Floyd, R. M., Hebert, P. D. N., Adams, A. M., Nagel, J., Girton, R., Newmaster, S. G., & Fenton, M. B. (2018). Approaches to integrating genetic data into ecological networks. Molecular Ecology, 28(2): 503–519. doi: 10.1111/mec.14941

Clark, N. J., & Clegg, S. M. (2017). Integrating phylogenetic and ecological distances reveals new insights into parasite host specificity. Mol. Ecol. 26, 3074–3086. https://doi.org/10.1111/mec.14101

de Vasconcelos, P., Falcão, L., Graciolli, G., & Borges, M. (2016). Parasite-host interactions of bat flies (Diptera: Hippoboscoidea) in Brazilian tropical dry forests. Parasitology Research, 115(1): 367–377. doi: 10.1007/s00436-015-4757-8

Dick, C. (2007). High host specificity of obligate ectoparasites Ecological Entomology, 32(5): 446–450. doi: 10.1111/j.1365-2311.2007.00836.x

Dick, C., & Dittmar, K. (2014). Parasitic Bat Flies (Diptera: Streblidae and Nycteribiidae): Host Specificity and Potential as Vectors. In: Klimpel S, Mehlhorn H (eds) Bats (Chiroptera) as Vectors of Diseases and Parasites: Facts and Myths. Springer Berlin Heidelberg, Berlin, Heidelberg, pp 131–155.

Dick, C. W., & Patterson, B. D. (2007). Against all odds: Explaining high host specificity in dispersal-prone parasites. International Journal for Parasitology, 37(8–9), 871–876. https://doi.org/10.1016/j.ijpara.2007.02.004

Dittmar, K., Dick, C. W., Patterson, B. D., Whiting, M. F., & Gruwell, M. E. (2009). Pupal Deposition and Ecology of Bat Flies (Diptera: Streblidae): Trichobius sp. (Caecus Group) in a Mexican Cave Habitat. Journal of Parasitology, 95(2), 308–314. https://doi.org/10.1645/GE-1664.1

Dobson, A. P. (1985). The population dynamics of competition between parasites. Parasitology, 91(2): 317–347. doi: 10.1017/S0031182000057401

Dormann, C.F., Gruber B. & Fruend, J. (2008). Introducing the bipartite Package: Analysing Ecological Networks. R news Vol 8/2, 8 – 11.

Folmer, O., Black, M., Hoeh, W., Lutz, R., & Vrijenhoek, R. (1994). DNA Primers for Amplification of Mitochondrial Cytochrome c Oxidase Subunit I from Diverse Metazoan Invertebrates. Molecular Marine Biology and Biotechnology 3 (5): 294–99.

Fox, J., & Weisberg, S. (2019). An R Companion to Applied Regression, Third edition. Sage, Thousand Oaks CA. https://socialsciences.mcmaster.ca/jfox/Books/Companion/.

Futuyma, D. J., & Moreno, G. (1988). The evolution of ecological specialization. Annual Review of Ecology and Systematics, 19(1), 207–233. https://doi.org/10.1146/annurev.es.19.110188.001231

Galen, S.C., Speer, K.A., & Perkins, S.L. (2019). Evolutionary lability of host associations promotes phylogenetic overdispersion of co-infecting blood parasites. J. Anim. Ecol. 88, 1936–1949. https://doi.org/10.1111/1365-2656.13089

Ge, X.-Y., Li, J.-L., Yang, X.-L., Chmura, A. A., Zhu, G., Epstein, J. H., Mazet, J. K., Hu, B., Zhang, W., Peng, C., Zhang, Y.-J., Luo, C.-M., Tan, B., Wang, N., Zhu, Y., Crameri, G., Zhang, S.-Y., Wang, L.-F., Daszak, P., & Shi, Z.-L. (2013). Isolation and characterization of a bat SARS-like coronavirus that uses the ACE2 receptor. Nature, 503(7477), 535–538. https://doi.org/10.1038/nature12711

Graciolli, G., & de Carvalho, C. J. B. (2001a). Moscas ectoparasitas (Diptera, Hippoboscoidea, Nycteribiidae) de morcegos (Mammalia, Chiroptera) do Estado do Paraná, Brasil. I. Basilia, taxonomia e chave pictórica para as espécies. Revista Brasileira de Zoologia, 18, 33–49. doi: 10.1590/S0101-81752001000500002

Graciolli, G., & de Carvalho, C. (2001b). Moscas ectoparasitas (Diptera, Hippoboscoidea) de morcegos (Mammalia, Chiroptera) do Estado do Paraná. II. Streblidae. Chave pictórica para gêneros e espécies. Revista Brasileira de Zoologia, 18(3): 907–960. doi: 10.1590/S0101-81752001000500002

Graciolli, G., & de Carvalho, C. J. B. (2012). Do fly parasites of bats and their hosts coevolve? Speciation in *Trichobius phyllostomae* group (Diptera, Streblidae) and their hosts (Chiroptera, Phyllostomidae) suggests that they do not. Revista Brasileira de Entomologia, 56(4), 436–450. https://doi.org/10.1590/S0085-56262012000400007

Hatcher, M., & Dunn, A. (2011). Parasites in ecological communities: from interactions to ecosystems. Cambridge University Press, Cambridge; New York

Hebert, P. D. N., Cywinska, A., Ball, S.L., & deWaard, J.R. (2003). Biological Identifications through DNA Barcodes. Proceedings of the Royal Society of London. Series B: Biological Sciences 270 (1512): 313–21. https://doi.org/10.1098/rspb.2002.2218

Hebert, P. D. N., Penton, E. H., Burns, J. M., Janzen, D. H., & Hallwachs, W. (2004). Ten species in one: DNA barcoding reveals cryptic species in the neotropical skipper butterfly Astraptes fulgerator. Proceedings of the National Academy of Sciences, 101(41), 14812– 14817. https://doi.org/10.1073/pnas.0406166101

Hiller, T., Brändel, S.D., Honner, B., Page, R.A., & Tschapka, M. (2020). Parasitization of bats by bat flies (Streblidae) in fragmented habitats. Biotropica 72, 617. https://doi.org/10.1111/btp.12757

Ingala, M., Simmons, N., & Perkins, S. (2018). Bats are an untapped system for understanding microbiome evolution in mammals. mSphere, 3(5): e00397–18. doi: 10.1128/mSphere.00397-18

Kembel, S. W., Cowan, P. D., Helmus, M. R., Cornwell, W. K., Morlon, H., Ackerly, D. D., Blomberg, S. P., & Webb, C. O. (2010). Picante: R tools for integrating phylogenies and ecology. Bioinformatics, 26(11), 1463–1464. https://doi.org/10.1093/bioinformatics/btq166.

Krasnov, B., Mouillot, D., Shenbrot, G., Khokhlova, I.S., & Poulin, R. (2011). Beta-specificity: The turnover of host species in space and another way to measure host specificity. International Journal for Parasitology, 41(1): 33–41. doi: 10.1016/j.ijpara.2010.06.001

Kunz, T. H., & Fenton, M. B. (2005). *Bat Ecology*. University of Chicago Press.

Lawton, J. H. (1999). Are there general laws in ecology? Oikos, 84(2), 177–192. JSTOR. https://doi.org/10.2307/3546712

Long, J. A. (2019). jtools: Analysis and Presentation of Social Scientific Data. R package version 2.0.0, https://cran.r-project.org/package=jtools

Marshall, A. (1981). The ecology of ectoparasitic insects. Academic Press Inc., (London) Ltd.

Massey, F. J. (1951). The Kolmogorov-Smirnov Test for Goodness of Fit. Journal of the American Statistical Association, 46(253), 68–78. doi: 10.2307/2280095

Medellín, R. A., Equihua, M., & Amin, M. A. (2000). Bat Diversity and Abundance as Indicators of Disturbance in Neotropical Rainforests. Conservation Biology, 14(6), 1666–1675. https://doi.org/10.1111/j.1523-1739.2000.99068.x

Oksanen J., Blanchet, F.G., Friendly, M., Kindt, R., Legendre, P., McGlinn, D., Minchin, P.R., O’Hara, R. B., Simpson, G.L., Solymos, P., Stevens, M.H.H., Szoecs, E., & Wagner, H. (2019). vegan: Community Ecology Package. R package version 2.5–6. https://CRAN.R/project.org/package=vegan

Olival, K. J., Dick, C. W., Simmons, N. B., Morales, J. C., Melnick, D. J., Dittmar, K., Perkins, S. L., Daszak, P., & DeSalle, R. (2013). Lack of population genetic structure and host specificity in the bat fly, *Cyclopodia horsfieldi*, across species of *Pteropus* bats in Southeast Asia. Parasites & Vectors, 6(1), 231. https://doi.org/10.1186/1756-3305-6-231

Pilosof, S., Dick, C. W., Korine, C., Patterson, B. D., & Krasnov, B. R. (2012). Effects of Anthropogenic Disturbance and Climate on Patterns of Bat Fly Parasitism. PLoS ONE, 7(7), e41487. https://doi.org/10.1371/journal.pone.0041487

Pinheiro, R. B. P., Félix, G. M. F., Chaves, A. V., Lacorte, G. A., Santos, F. R., Braga, É. M., & Mello, M. A. R. (2016). Trade-offs and resource breadth processes as drivers of performance and specificity in a host–parasite system: a new integrative hypothesis. International Journal for Parasitology, 46(2), 115–121. https://doi.org/10.1016/j.ijpara.2015.10.002

Poulin, R. (2007). Are there general laws in parasite ecology? Parasitology, 134(6), 763–776. https://doi.org/10.1017/S0031182006002150

Poulin, R., Krasnov, B., Mouillot, D. (2011). Host specificity in phylogenetic and geographic space. Trends in Parasitology, 27(8): 355–361. doi: 1016/j.pt.2011.05.003

Presley, S. (2011). Interspecific aggregation of ectoparasites on bats: importance of hosts as habitats supersedes interspecific interactions. Oikos 120(6): 832–841. doi: 10.1111/j.1600-0706.2010.19199.x

R Core Team (2020). R: A language and environment for statistical computing. R Foundation for Statistical Computing, Vienna, Austria. https://www.R-project.org/

Reed, D. L., & Hafner, M. S. (1997). Host Specificity of Chewing Lice on Pocket Gophers: A Potential Mechanism for Cospeciation. Journal of Mammalogy, 78(2), 655–660. https://doi.org/10.2307/1382916

Rivera-García, K., Sandoval-Ruiz, C., Saldaña-Vázquez, R., & Schondube, J. (2017). The effects of seasonality on host–bat fly ecological networks in a temperate mountain cave. Parasitology, 144(5): 692–697. doi: 10.1017/S0031182016002390

Rojas, D., Vale, Á., Ferrero, V., & Navarro, L. (2011). When did plants become important to leaf-nosed bats? Diversification of feeding habits in the family Phyllostomidae: Evolution of feeding habits in Phyllostomid bats. Molecular Ecology, 20(10): 2217–2228. doi: 10.1111/j.1365-294X.2011.05082.x

Shi, J., & Rabosky, D. (2015). Speciation dynamics during the global radiation of extant bats. Evolution, 69(6): 1528–1545. doi: 10.1111/evo.12681

Tello, J. S., Stevens, R. D., & Dick, C. W. (2008). Patterns of species co-occurrence and density compensation: a test for interspecific competition in bat ectoparasite infracommunities. Oikos, 117(5), 693–702. https://doi.org/10.1111/j.0030-1299.2008.16212.x

ter Hofstede, H., Fenton, M., Whitaker Jr., J. (2004). Host and host-site specificity of bat flies (Diptera: Streblidae and Nycteribiidae) on Neotropical bats (Chiroptera). Canadian Journal of Zoology, 82(4): 616–626. doi: 10.1139/z04-030

Vázquez, D. P., & Simberloff, D. (2002). Ecological Specialization and Susceptibility to Disturbance: Conjectures and Refutations. The American Naturalist, 159(6), 606–623. https://doi.org/10.1086/339991

Voss, R. S., Fleck, D. W., Strauss, R. E., Velazco, P. M., & Simmons, N. B. (2016). Roosting Ecology of Amazonian Bats: Evidence for Guild Structure in Hyperdiverse Mammalian Communities. American Museum Novitates, (3870), 1–43. doi:10.1206/3870.1

Wenzel, R. (1976). The streblid batflies of Venezuela (Diptera: Streblidae). *Brigham Young University Science Bulletin*, Biological Series, 20(4): 1.

Zarazúa-Carbajal M., Saldaña-Vázquez R., Sandoval-Ruiz, C., Stoner, K. E., & Benitez-Malvido, J. (2016). The specificity of host-bat fly interaction networks across vegetation and seasonal variation. Parasitology Research, 115(10): 4037–4044. doi: 10.1007/s00436-016-5176-1

Zhou, P., Yang, X. L., Wang, X. G., Hu, B., Zhang, L., Zhang, W., … & Shi, Z.L. (2020). A pneumonia outbreak associated with a new coronavirus of probable bat origin. Nature, 579 (7798), 270–273. https://doi.org/10.1038/s41586-020-2012-7

