## Supplementary Figure 1 for "Phylogenetic and Ecological Trends in Specialization: Disentangling the Drivers of Ectoparasite Host Specificity"

**Supplementary Figure 1***.* Map of study site within the Atlantic Forest area of the eastern Brazil coast (left) and the 13 sampling fragments included in this study (inset, right). Three sites (REGUA 1, REGUA 2, and REGUA 3) were located within the bounds of the Reserva Ecológica de Guapiaçu, whereas the remaining 10 sites were located within forest fragments on the outskirts of the reserve. Green polygons represent forested area, grey indicates the surrounding matrix.

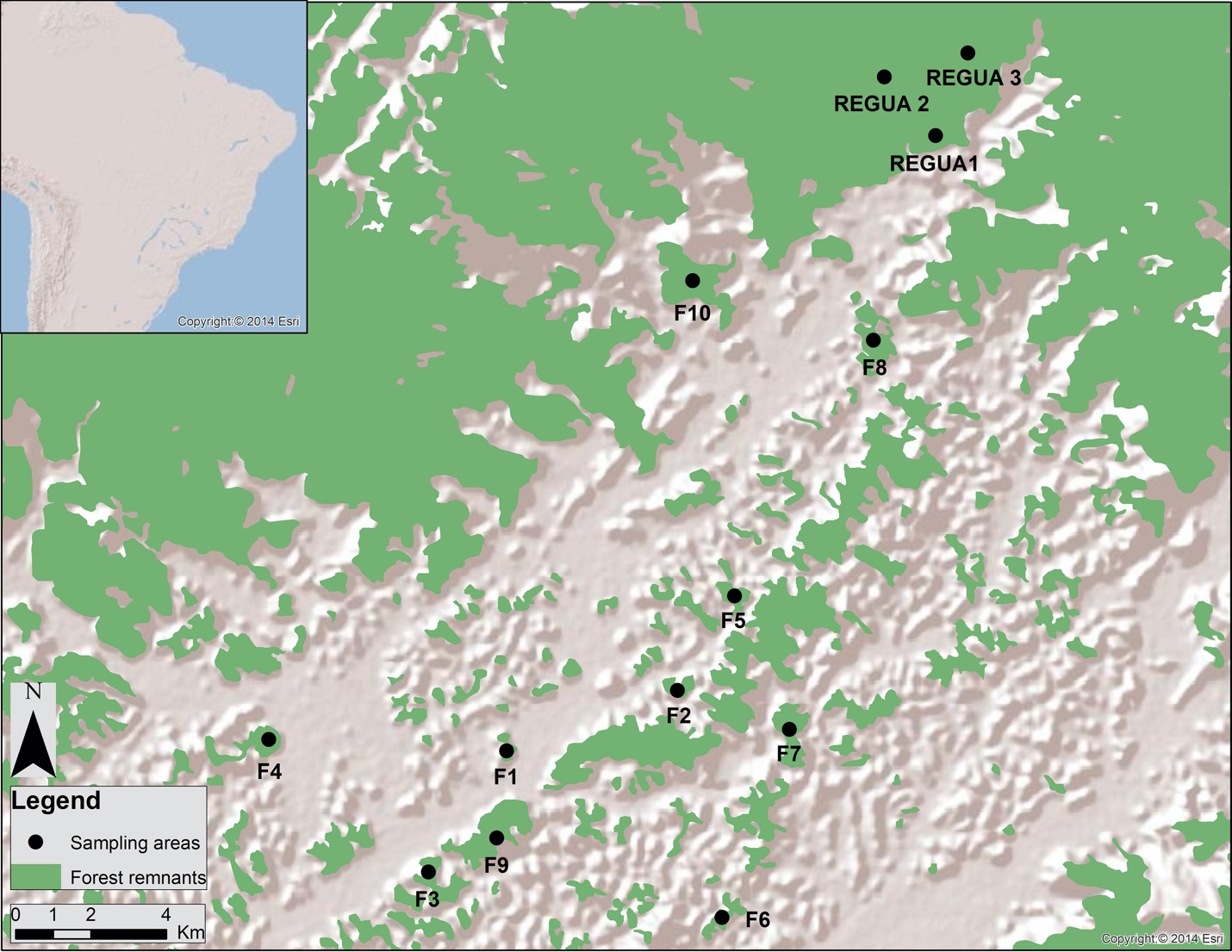
