## Supplement 2 for "Phylogenetic and Ecological Trends in Specialization: Disentangling the Drivers of Ectoparasite Host Specificity"

**Supplement 2: DNA Extraction and Amplification Details/Protocols**

1. Extraction

We extracted genomic DNA from bat flies using DNeasy Tissue Extraction Kit (QIAGEN Inc), or Zymo Research ZymoBIOMICS DNA Miniprep Kit (Zymo Research, Irvine, CA). Because individual bat flies differed in their physical condition (some were crushed by forceps while others remained entirely intact), we processed “damaged” and “intact” flies via two different extraction protocols. DNA from all “damaged” flies (approximately 275 individuals) was extracted using the QIAGEN kit, whereas the “intact” flies were split between QIAGEN and Zymo kits for no reason other than timing of the study; laboratory-provided QIAGEN kits became unavailable to us towards the end of our DNA extraction period, and Zymo kits were ordered as a replacement. We have no reason to believe the use of these different extraction kits had any effect on the quality of DNA extraction or amplification of targeted genes.

a) **“Damaged” flies**: Individual flies were removed from their ethanol tubes and allowed to air dry for approximately 10 minutes. The DNA extraction followed the standard QIAGEN protocol with the following modifications:

i) Proteinase K digestion was conducted in an incubator at 56°C on a rotating	mixing plate set to 350rpm. Digestions continued overnight for at least 24 hours.

ii) Elution was performed by adding 100 $\mu$L of 56°C incubated distilled H_2_0, and	then re-eluting the 100$\mu$L flow-through. This was done to maximize DNA yield.

b) **“Intact” flies:** To facilitate the digestion step of extraction, “intact” flies were transferred to individual microbead-filled tubes and submerged in 50$\mu$L of 1X PBS buffer solution. Tubes were placed on a Disruptor Genie (Scientific Industries, Inc.) and underwent 3 homogenization intervals of 30 seconds each, with a 60-second rest period in between each interval. Following homogenization, DNA was extracted either via QIAGEN protocol (at beginning of the study, with a 24-hour Proteinase-K digestion period, conducted in an incubator at 56°C on a rotating mixing plate set to 500 rpm) or Zymo Quick-gDNA protocol (end of study, no Proteinase-K digestion). Elution for both QIAGEN and Zymo protocols was performed by adding 100 $\mu$L of 56°C incubated distilled H_2_0, and then re-eluting the 100$\mu$L flow-through.

Following DNA extraction, DNA was either processed immediately for PCR amplification, or stored in a -80°C laboratory freezer for PCR to be performed at a later date.

2. PCR

We amplified a 645-basepair region of the cytochrome oxidase-I (COI) mitochondrial gene using forward primer LC01490 (5’- GGTCAACAAATCATAAAGATATTGG-3’) and reverse primer HC02198 (5’-TAAACTTCAGGGTGACCAAAAAAT-3’) (Folmer et al. 1994) and the following PCR protocol: 1-minute denaturation at 94°C followed by 5 cycles of 94°C for 40 s, 45°C for 40 s, 72°C for 1 min and 35 cycles of 94°C for 40 s, 51°C for 40 s, 72°C for 1 min, and 72°C for 5 min. Master mix concentrations were as follows: 757 $\mu$L TopTaq, 151.5 $\mu$L Coral Dye, 15.2 $\mu$L LCO1490 forward primer, 15.2 $\mu$L HCO2190 reverse primer, and 474.7 $\mu$L H_2_O. 1 $\mu$L of template DNA was added to 14 $\mu$L of master mix for a 15 $\mu$L total PCR reaction.

We amplified a 742-baspair region of the carbamoyl-phosphate synthetase (CAD) nuclear gene using a nested primer method as described in Petersen et al. (2007) with forward primer 787F (Round 1: GGD GTN ACN ACN GCN TGY TTY GAR CC; Round 2: GTN GTN AAR ATG CCN MGN TGG GA) and reverse primer 1124R (Round 1: CAT NCG NGA RAA YTT RAA RCG ATT YTC; Round 2: TTN GGN AGY TGN CCN CCC AT) and the following PCR protocol: 1-minute denaturation at 94°C followed by 5 cycles of 94°C for 40 s, 45°C for 40 s, 72°C for 1 min and 35 cycles of 94°C for 40 s, 51°C for 40 s, 72°C for 1 min, and 72°C for 5 min. Master mix concentrations for both nested rounds were as follows: 612 $\mu$L TopTaq, 102 $\mu$L Coral Dye, 51 $\mu$L forward and 51 $\mu$L reverse primer, and 408 $\mu$L H_2_0. 1 $\mu$L of template DNA was added to 24 $\mu$L of master mix for a 25 $\mu$L total PCR reaction.

We visualized PCR products on 1.5% agarose gel through electrophoresis and cleaned PCR amplicons using AMPure XP beads (Beckman Coulter, Inc.). We then cycle sequenced cleaned PCR products in 10μL reactions using Big Dye Terminator v3.1 (Life Technologies Corporation, Carlsbad, CA) chemistry following the standardly used protocol. CycleSeq master mix concentrations were as follows: 99 $\mu$L Big Dye, 99$\mu$L of 1$\mu$M diluted primer, 99 $\mu$L EB and 495 $\mu$L H_2_O. 2 $\mu$L of cleaned PCR product was added to 8 $\mu$L of master mix for a 10 $\mu$L total sequencing reaction. Reactions were cleaned using AMPure XP beads and then sequenced on a 3730xl Applied Biosystems (ABI) machine (Thermo Fisher Scientific).
